# Epo-IGF1R crosstalk expands stress-specific progenitors in regenerative erythropoiesis and myeloproliferative neoplasm

**DOI:** 10.1101/2022.06.27.497649

**Authors:** Hsi-Hsien Hsieh, Huiyu Yao, Yue Ma, Yuannyu Zhang, Xue Xiao, Helen Stephens, Stephen S. Chung, Lin Xu, Jian Xu, Raajit K. Rampal, Lily Jun-shen Huang

**Author notes:** Correspondence: Lily Huang, Department of Cell Biology, University of Texas Southwestern Medical Center, Dallas, TX 75390-9039;. Authorship note: HH and HY contributed equally to this work.

## Abstract

We find that in regenerative erythropoiesis, the erythroid progenitor landscape is reshaped, and a previously undescribed progenitor population with CFU-E activity (stress CFU-E/sCFU-E) is markedly expanded to restore the erythron. sCFU-E are targets of erythropoietin (Epo) and sCFU-E expansion requires signaling from the Epo receptor (EpoR) cytoplasmic tyrosines. Molecularly, Epo promotes sCFU-E expansion via JAK2/STAT5-dependent expression of IRS2, thus engaging the pro-growth signaling from the IGF1 receptor (IGF1R). Inhibition of IGF1R/IRS2 signaling impairs sCFU-E cell growth, whereas exogenous IRS2 expression rescues cell growth in sCFU-E expressing truncated EpoR lacking cytoplasmic tyrosines. This sCFU-E pathway is the major pathway involved in erythrocytosis driven by the oncogenic JAK2 mutant, JAK2(V617F), in myeloproliferative neoplasm. Inability to expand sCFU-E cells by truncated EpoR protects against JAK2(V617F)-driven erythrocytosis. In myeloproliferative neoplasm patient samples, the number of sCFU-E like cells increases, and inhibition of IGR1R/IRS2 signaling blocks Epo-hypersensitive erythroid cell colony formation. In summary, we identify a new stress-specific erythroid progenitor cell population that links regenerative erythropoiesis to pathogenic erythrocytosis.

**Key Points:**

- Epo-induced IRS2 allows engagement of IGF1R signaling to expand a previously unrecognized progenitor population in erythropoietic stress.
- Truncated EpoR does not support stress CFU-E expansion and protects against JAK2(V617F)-driven erythrocytosis in MPN.

## INTRODUCTION

Hematologic homeostasis in adult humans requires the production of roughly 200 billion new erythrocytes per day. Erythropoiesis is a multistep process that begins when hematopoietic stem cells differentiate into erythroid progenitors. The earliest committed erythroid progenitors are the burst-forming unit-erythroid (BFU-E), which divide and differentiate into colony-forming unit-erythroid (CFU-E). CFU-E progenitors further proliferate and differentiate into erythroid precursors, which undergo terminal maturation to generate erythrocytes.

Erythropoietin (Epo) is the principal cytokine that controls erythropoiesis.^1-3^ Epo binding to Epo receptor (EpoR) activates the tyrosine kinase JAK2 which associates with the EpoR cytoplasmic domain. Activated JAK2 phosphorylates many of the tyrosine residues in the EpoR cytoplasmic domain, leading to docking of signaling proteins and the subsequent activation of the STAT5, PI3K/Akt, and MAPK pathways.^4-6^ Together, these pathways promote erythroid cell survival, proliferation, and differentiation.

At steady-state, normal Epo levels support basal erythropoiesis to replace the clearance of aged erythrocytes. Erythropoietic stress such as bleeding causes Epo levels to surge, dramatically increasing erythropoiesis via a process termed stress erythropoiesis. Stress erythropoiesis is critical for the recovery and survival from blood loss, anemia of multiple etiologies or from therapeutic procedures such as chemotherapy and stem-cell transplantation. Despite the importance of stress erythropoiesis, our understanding of this process remains incomplete.

Stress erythropoiesis differs from basal erythropoiesis in several ways. Basal erythropoiesis is achieved by fine-tuning survival and proliferation of erythroid precursors downstream of CFU-E, whereas stress erythropoiesis expands both erythroid precursors and progenitors.^7-10^ While both basal and stress erythropoiesis are each regulated by EpoR, only stress erythropoiesis requires an intact EpoR cytoplasmic domain. Mice expressing a truncated EpoR lacking all cytoplasmic domain tyrosines has a near-normal basal hematocrit but is deficient in its response to stress.^11,12^ In addition to Epo, stress erythropoiesis also involves corticosteroids, stem cell factor (SCF), and signaling from bone morphogenetic protein 4 (BMP4).^13-15^

Myeloproliferative neoplasms (MPN) are a group of chronic myeloid malignancies characterized by clonal expansion of one or more myelo-erythroid lineage cells. Clinically, MPNs present as overproduction of erythrocytes (polycythemia vera, PV) or platelets (essential thrombocytosis, ET), or as bone marrow fibrosis (primary myelofibrosis, PMF). MPNs can transform into acute myeloid leukemias, which commonly have poor prognosis.^16-18^ In PV, erythropoiesis progresses at an aberrantly high rate even in the absence of increased Epo due to somatic mutations (most commonly V617F) in JAK2 that constitutively activates JAK2 kinase activity.^19,20^ While Epo-independent, JAK2(V617F)-induced erythrocytosis still requires the EpoR to engage downstream signaling proteins.^21^

Although EpoR signaling in more differentiated erythroblasts has been well characterized,^22-24^ EpoR signaling in earlier progenitors is less well understood. In this study, we discovered a new population of progenitors that are able to form CFU-E like colonies, here termed stress CFU-E (sCFU-E), that are specifically expanded by erythropoietic stress. Failure to stimulate sCFU-E expansion in mice expressing truncated EpoR blocks both regenerative erythropoiesis and JAK2(V617F)-driven erythrocytosis, suggesting that oncogenic JAK2 hijacks sCFU-E to promote erythrocytosis in MPN.

## METHODS

### Mice, phlebotomy, Epo and phenylhydrazine injection

3-6 month-old mice were used for all experiments. Phlebotomy was performed by submandibular bleeding (400μL) followed by fluid replacement with normal saline twice over 6 hrs. For phenylhydrazine (PHZ) treatments, mice were injected intraperitoneally with 62.5 mg/kg (low dose) or 87.5 mg/kg (high dose) PHZ on day 0 and day 1. For Epo injections, 100 U of Epoetin alpha (Amgen) was injected once subcutaneously.

### Flow cytometry

For progenitor analyses, BFU-E and CFU-E progenitors were identified as described.^25^ Data were acquired on a LSRII, Fortessa, or Aria (BD Biosciences) flow cytometer and analyzed with FlowJo software (Tree Star, CA).

### *In vitro* culture of erythroid progenitors

BFU-E, sCFU-E and CFU-E were isolated from murine bone marrow by flow cytometry sorting after immune-depletion of lineage (Lin)-committed cells and hematopoietic stem cells (Sca1) using biotin-conjugated antibodies followed by streptavidin-conjugated magnetic resin. Sorted cells were subsequently cultured in StemPro34 media supplemented with 2 U/ml Epo, 100 ng/ml SCF, 40 ng/ml IGF1, and 1 μM dexamethasone. Cells were analyzed at indicated time.

### Human MPN patient samples

Bone marrow mononuclear samples were obtained from consented and deidentified PV, lymphoma and MGUS patients. Human BFU-E and CFU-E were identified as described.^26^ For *in vitro* cultures, sorted PV CD34^+^CD36^-^ cells were first expanded in StemSpan SFEM media with StemSpan CC100 (both from StemCell Technologies) for 3 days. Subsequently, cells were cultured in SFEM media with CC100 and Epo (3U/ml) (day 0). As controls, normal CD34^+^ cells (Cooperative Center of Excellence in Hematology at Fred Hutch) were sorted, expanded, and cultured in parallel.

### Statistical analyses

Data are reported as means ± standard deviation (SD). Statistical significance was determined by using Student t-test or analysis of variance (ANOVA). A value of P<0.05 was considered statistically significant. Normality tests and statistical analyses were performed using Prism (Graphpad).

### Study approval

Deidentified human samples were acquired through the Hematologic Malignancies Tissue Bank at UT Southwestern under institutional IRB with informed consent of all participants. All mouse studies were approved by institutional IACUC.

### Additional methods in supplement

Detailed methods are in Supplemental Methods.

## RESULTS

### Phlebotomy induces expansion of a new population of erythroid progenitors

To examine early erythroid progenitors, we used a method that allows flow cytometric identification of murine BFU-E and CFU-E in adult hematopoietic tissues.^25^ In this assay, Lin^-^Kit^+^CD55^+^CD105^+^ cells are dissected into BFU-E (CD150^+^CD71^-^) and CFU-E (CD150^-^CD71^+^) (Fig. 1A). Interestingly, a drastic expansion of an intermediate cell population that was CD150^+^CD71^+^ was observed upon phlebotomy, a physiological erythropoietic stress (Fig. 1B). We designated this population as “sCFU-E” for “stress CFU-E” based on our subsequent characterization. During erythron recovery, wild-type mice exhibited a nadir of RBC number 2 days following phlebotomy and recovered a normal hematocrit by day 9 (Fig. 1C). Expansion of sCFU-E cells was observed one day after phlebotomy and coincided with a decrease in BFU-E. Thereafter, the number of sCFU-E cells decreased while and CFU-E cells increased, followed by an increase in Ter119^+^ erythroblasts (Fig. 1D-G). Similar observations were made in the spleen (Fig. 1D-G). These observations suggest that sCFU-E are involved in regenerative erythropoiesis.

**Fig. 1.**
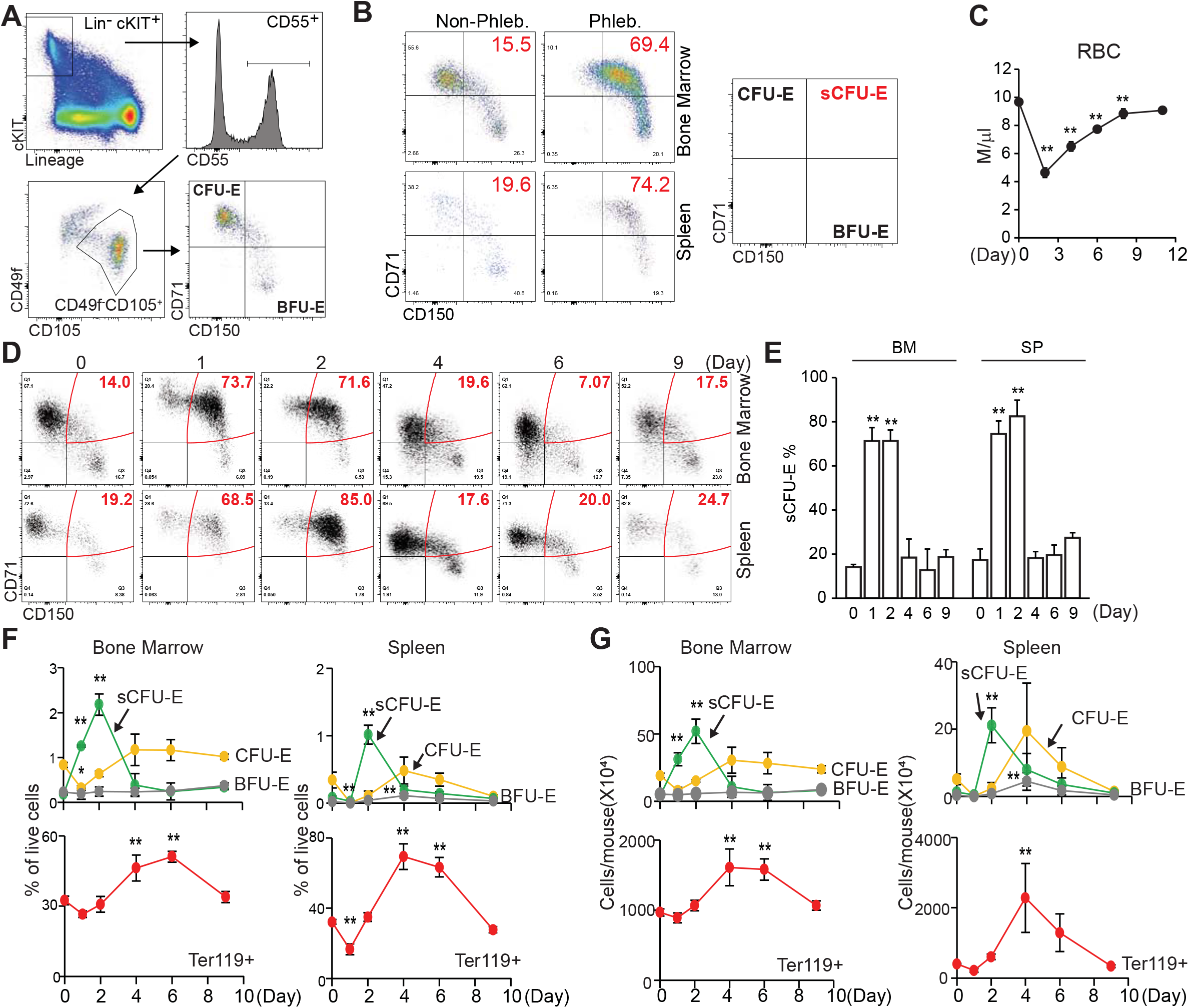
sCFU-E cells expand in erythropoietic stress. (A) Flow cytometric gating strategy. (B) Percentages of sCFU-E cells increase in the bone marrow and spleen of phlebotomized mice 2 days post phlebotomy (C) Red blood cell (RBC) counts on indicated day post phlebotomy. (D) Representative flow cytometry plots of temporal sCFU-E increases in the bone marrow and spleen of phlebotomized mice. (E) Quantification of percentage changes of sCFU-E in (D). (F) Quantification of BFU-E, sCFU-E, CFU-E and Ter119^+^ cell percentages in phlebotomized mice at indicated times. (G) Total numbers of BFU-E, sCFU-E, CFU-E and Ter119^+^ cells in phlebotomized mice at indicated times. Data represent the mean ± SD. *P < 0.05, **P < 0.01 analyzed by one-way ANOVA.

To examine the temporal relationship between these progenitors, *in vitr*o cultures were set up, where BFU-E cells from normal or phlebotomized mice were prospectively isolated and cultured in media containing SCF, Epo and dexamethasone to mimic stress erythropoiesis. Consistent with results *in vivo*, BFU-E gave rise to sCFU-E in 24 hours, which further differentiate into CFU-E (Supplemental Fig. 1). Therefore, as erythropoiesis proceeds as a continuum,^25,27,28^ the temporal order of development proceeds from BFU-E to sCFU-E to CFU-E.

### Characterization of sCFU-E cells

BFU-E, sCFU-E and CFU-E from bone marrow of phlebotomized mice were isolated and analyzed by histology and colony assays. By Giemsa staining, sCFU-E were more similar to CFU-E and were larger than BFU-E, consistent with their higher forward scatter intensity (Fig. 2A-B). sCFU-E and CFU-E also had more prominent nucleoli as compared to BFU-E (Fig. 2A). In colony assays, sCFU-E generated uni-focal CFU-E-like colonies by day 2, not large multi-focal “burst” colonies like BFU-E which usually take 5-7 days (Fig. 2C). That both sCFU-E and CFU-E formed CFU-E like colonies is consistent with our observation that the predominant colonies expanded following phlebotomy were CFU-E but not BFU-E colonies (Supplemental Fig. 2). Interestingly, sCFU-E colonies were significantly larger and had about 3 times as many cells as CFU-E colonies (Fig. 2C-D), suggesting that they possess higher proliferative potential. Indeed, sorted sCFU-E proliferated longer and generated more erythroid progeny compared to CFU-E *in vitro* (Fig. 2E). Based on the morphology of colonies they made, and the fact that sCFU-E cells are specifically up-regulated in stress, we have named them “stress CFU-E” or “sCFU-E”.

**Fig. 2.**
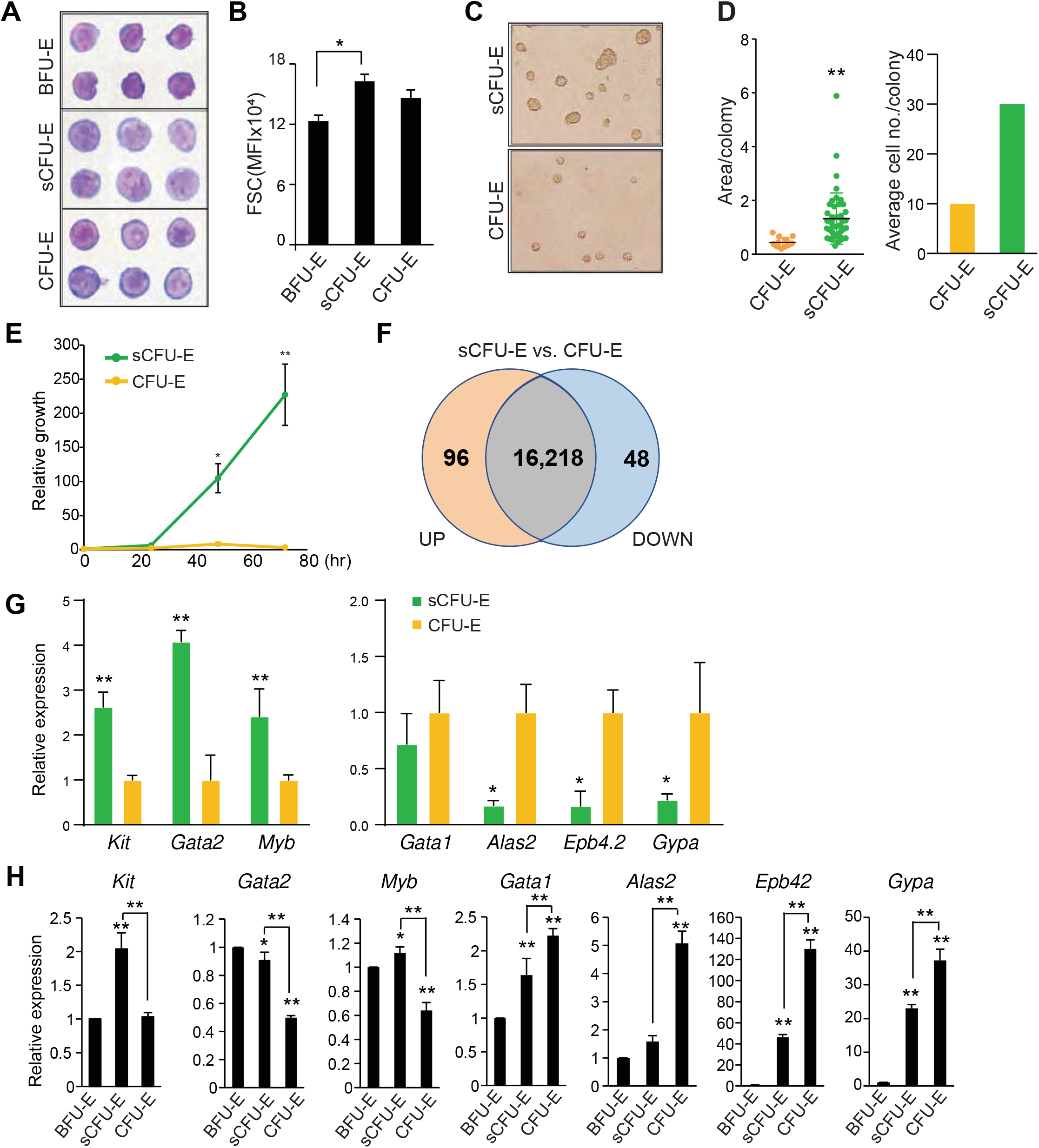
sCFU-E exhibit higher growth potential and express lower levels of erythroid committed genes compared to CFU-E. (A) Histology staining of sorted BFU-E, sCFU-E and CFU-E. (B) Quantification of forward scatter (FSC) median fluorescence intensity, an indicator of cell size, by flow cytometry (C) sCFU-E cells generate uni-focal colonies on day 2. Colonies generated from sCFU-E are larger than those from CFU-E cells. (D) Quantification of area per colony and average cell number per colony in (C). (E) Sorted sCFU-E cells generate more progenies than CFU-E *in vitro*. (F) Comparison of sCFU-E and CFU-E transcriptome by RNA-seq. The number of genes with expression greater (up) or less (down) than 1.5-fold are indicated. (G) Relative expression of indicated genes in sCFU-E vs. CFU-E cells from RNA-seq data. (H) Expression of indicated genes in sorted BFU-E, sCFU-E and CFU-E by qPCR. Gene expression is first normalized to β-actin and then to expression in BFU-E. Significant differences between sCFU-E and CFU-E are specified. Data represent the mean ± SD. Statistically significant differences indicated on top of each bar are in comparison to BFU-E. *P < 0.05, **P < 0.01 analyzed by Student’s t-test or one-way ANOVA.

To compare sCFU-E and CFU-E, gene expression across the transcriptome were analyzed by RNA-seq using triplicate biological samples sorted from phlebotomized mouse marrow. We found that transcriptomes of sCFU-E and CFU-E were similar, with only 118 genes differentially expressed (fold change >1.5 and P<0.05, Fig. 2F). While no pathways were enriched in the differentially expressed genes (data not shown), several genes associated with an earlier stem/progenitor signature were enriched in sCFU-E and genes associated with committed erythropoietic functions were enriched in CFU-E (Fig. 2G). For example, the transcription factors Gata2 and Myb, as well as the receptor for stem cell factor, c-Kit, were higher in sCFU-E, whereas *Alas2* (erythroid specific delta-aminolevulinate synthase 2), *Epb4*.*2* (erythrocyte membrane protein band 4.2), and *Gypa* (glycophorin A) were expressed at higher levels in CFU-E (Fig. 2G). RNAseq results were corroborated with quantitative PCR analyses using sorted BFU-E, sCFU-E and CFU-E from phlebotomized mice (Fig. 2H). These results suggest that in erythropoietic stress, an unexpected shift occurs within the erythroid progenitor compartment, where a new cell population with higher proliferation potential, sCFU-E, is disproportionally expanded relative to CFU-E, increasing erythropoietic output.

### sCFU-E expansion requires distal EpoR signaling

Because Epo critically regulates stress erythropoiesis, we tested whether sCFU-E are direct targets of Epo by analyzing erythroid progenitors following Epo injection. sCFU-E significantly expanded in both marrow and spleen starting as early as 6 hours post Epo injection (Fig. 3B-C). We also examined sCFU-E expansion in a knock-in mouse model expressing a truncated EpoR lacking all cytoplasmic tyrosine residues (EpoR-HM).^12^ This mouse was shown be viable, but can only support basal erythropoiesis, not stress erythropoiesis (Fig. 3A). ^3,11,12^ Because this truncated EpoR represents a “core minimal receptor” sufficient for basal erythropoiesis, we refer to it as EpoR(core) herein (Fig. 3A). Contrary to what was observed in wild-type mice, Epo injection failed to increase sCFU-E in EpoR(core) mice (Fig 3B-C). EpoR(core) mice also failed to expand sCFU-E in response to phlebotomy (Fig. 3D) and had a slower recovery (Fig. 3E). EpoR(core) mice also recovered slower and succumbed to hemolysis challenge induced by high dose phenylhydrazine (Fig. 3F-G).

**Fig. 3.**
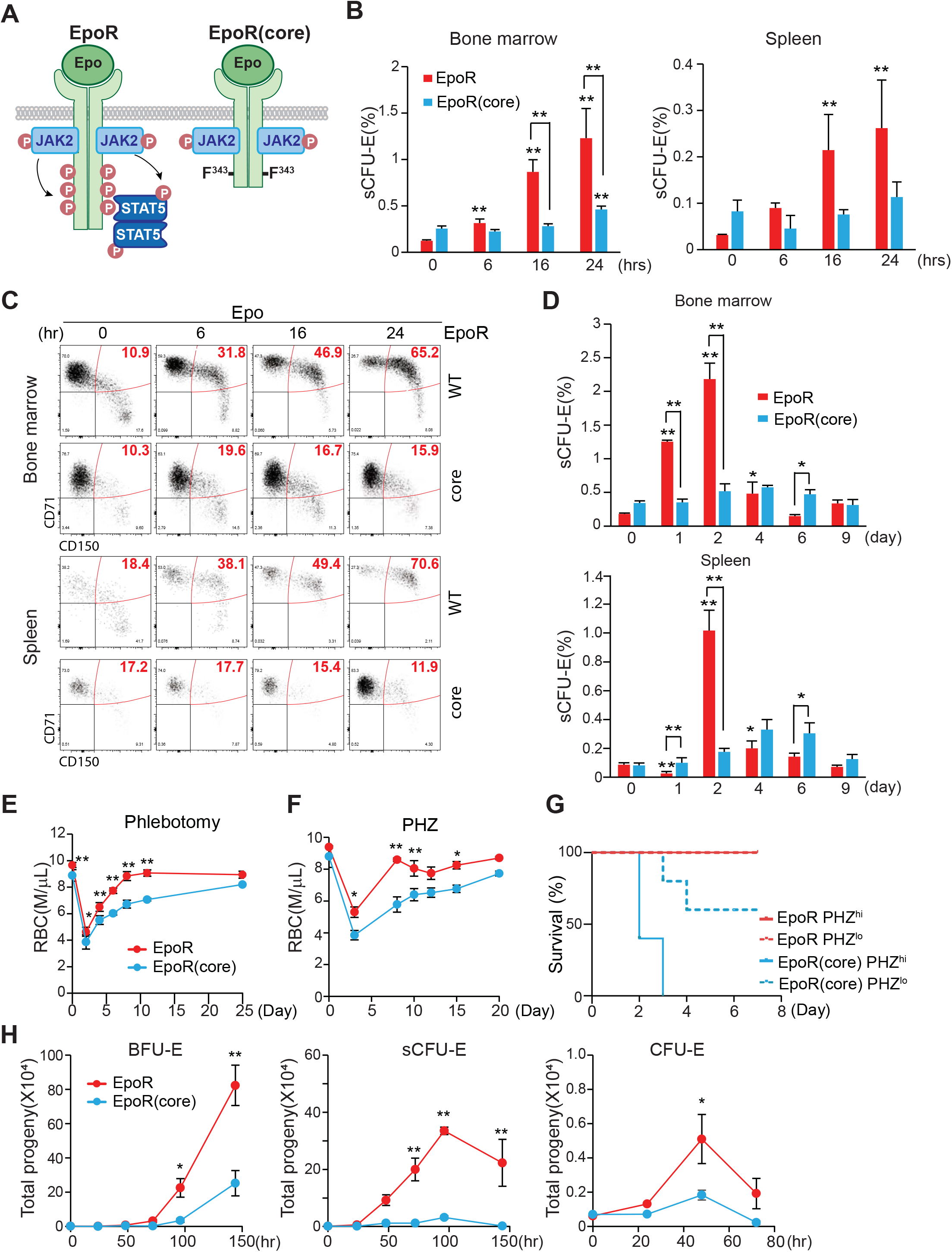
sCFU-E expansion is impaired in mice expressing truncated EpoR. (A) Diagrams of full-length EpoR and EpoR(core). (B) Epo-induced sCFU-E expansion is defective in EpoR(core) mice. Percentages of sCFU-E cells in the bone marrow or spleen were quantified at indicated times post Epo injection in mice expressing wild-type EpoR or EpoR(core). Statistically significant differences indicated on top of each bar are in comparison to time 0, whereas significant differences between EpoR and EpoR(core) at specific time points are specified. (C) Representative flow cytometry data from (B). (D) sCFU-E expansion post phlebotomy is defective in EpoR(core) mice. Statistically significant differences indicated on top of each bar are comparison with day 0, whereas significant differences between EpoR and EpoR(core) at specific time points are specified. (E) RBC counts post phlebotomy in mice expressing EpoR or EpoR(core) at indicated times. (F) RBC counts post phenylhydrazine (PHZ)-induced hemolysis in mice expressing EpoR or EpoR(core) at indicated times. (G) EpoR(core) mice succumb to erythropoietic stress elicited by PHZ treatment. Dosing of 62.5 mg/kg (PHZ^lo^) or 87.5 mg/kg (PHZ^hi^) is as indicated. (H) Sorted BFU-E and sCFU-E from EpoR(core) mice generated dramatically less erythroid progenies. Ter119^+^ erythroid progenies are enumerated at indicated time harvested from *in vitro* culture. Data represent the mean ± SD. *P < 0.05, **P < 0.01 analyzed by two-way ANOVA.

Consistent with the impaired sCFU-E expansion in EpoR(core) mice, sorted sCFU-E from EpoR(core) mice generated significantly fewer progeny compared to those from wild-type mice *in vitro* (Fig. 3H). Similarly, the upstream BFU-E from EpoR(core) generated less progenies. Sorted CFU-E from EpoR(core) mice also generated fewer progenies, but the degree of reduction was mild. Ter119^+^ erythroblasts, especially FSC^large^CD71^hi^ erythroblasts, expanded normally in EpoR(core) mice (Supplemental Fig. 3).^29^ These results suggest despite the ability to increase late erythroblasts, failure of sCFU-E expansion restricted erythroid output of EpoR(core) mice in stress. Moreover, EpoR cytoplasmic tyrosines are necessary for sCFU-E expansion.

### The STAT5 pathway is necessary and sufficient for sCFU-E expansion

To probe the underlying mechanism of sCFU-E expansion, we compared proliferation and apoptosis in sCFU-E in phlebotomized vs. normal mice. In the marrow, sCFU-E showed higher proliferation in phlebotomized mice compared to non-phlebotomized mice. In the spleen, both higher proliferation and reduced apoptosis was observed (Fig. 4A-B). Therefore, sCFU-E expansion involves enhanced proliferation and reduced apoptosis in erythropoietic stress.

**Fig. 4.**
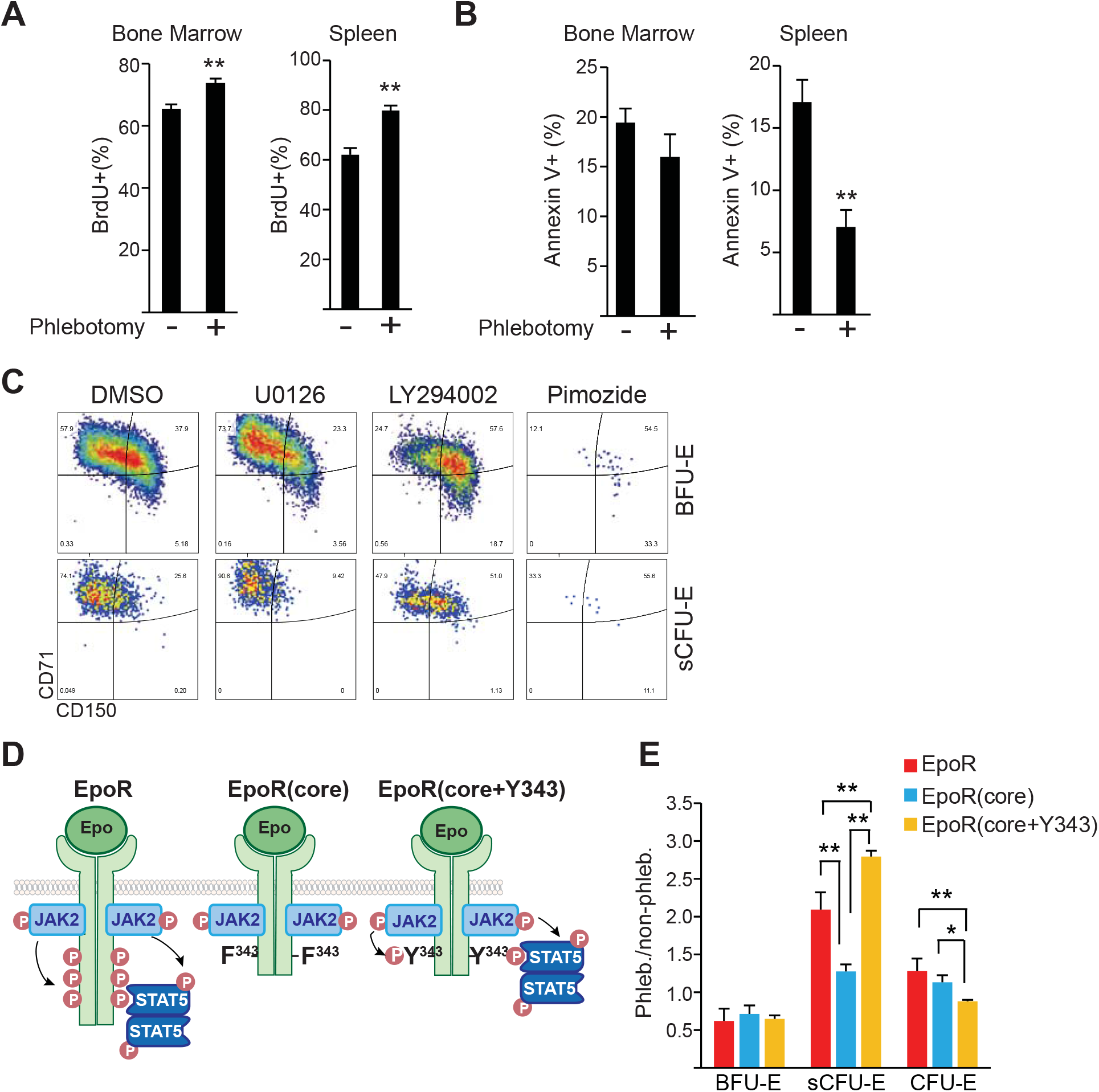
STAT5 signaling is essential for sCFU-E growth. (A) Proliferation increases in sCFU-E cells post phlebotomy. (B) Apoptosis decreases in sCFU-E cells post phlebotomy. (C) Inhibitors to STAT5 abolish sCFU-E growth. Sorted BFU-E and sCFU-E cells are cultured in 10μM inhibitors or vehicle control (DMSO) and analyzed after 48 hours. (D) Diagrams of EpoR(core) and EpoR(core+Y343). (E) Y343 in EpoR rescues STAT5 binding and sCFU-E expansion in EpoR(core+Y343) mice. Data represent the mean ± SD. *P < 0.05, **P < 0.01 analyzed by Student’s t-test or two-way ANOVA.

Three major pathways downstream of EpoR are the STAT5, PI3K/Akt, and MAPK pathways. We treated sorted BFU-E and sCFU-E *in vitro* with pathway inhibitors and examined the effects. STAT5 inhibitor pimozide completely abolished sCFU-E cell growth, while PI3K and MAPK inhibitors had much weaker effects (Fig. 4C). This is consistent with prior findings that STAT5 deficient mice have a near-normal hematocrit but are deficient in erythropoietic stress response.^10^

To verify the role of STAT5 in sCFU-E expansion, we used another murine model, EpoR-H (here refer to as EpoR(core+Y343)), that expresses EpoR(core) with Tyr343 restored (Fig. 4D).^12^ EpoR Tyr343 is a major STAT5 binding site and rescues both STAT5 activation and erythropoietic stress response in EpoR(core).^11^ EpoR(core+Y343) rescued sCFU-E expansion upon phlebotomy (Fig. 4E), suggesting that the EpoR/JAK2/STAT5 signaling pathway promotes sCFU-E expansion in stress erythropoiesis.

### The IGF1R/IRS2 pathway regulates sCFU-E expansion

We examined several known STAT5 target genes downstream of Epo in sCFU-E cells, including *Cish*, a feedback negative regulator of cytokine receptor signaling, *Bcl2l1*, an anti-apoptosis modulator, and *Fam132b* (also known as erythroferron), which regulates iron distribution. While these targets were all induced in CFU-E, they were hardly induced in sCFU-E (Fig. 5A). In contrast, *Irs2* (insulin receptor substrate-2) was acutely induced in sCFU-E by Epo (Fig. 5A) or phlebotomy (Fig. 5B). Consistent with the defect of EpoR(core) but not EpoR(core+Y343) to support sCFU-E expansion, Epo-induced *Irs2* expression was hampered in EpoR(core) and was restored in EpoR(core+Y343) sCFU-E (Fig. 5C-D).

**Fig. 5.**
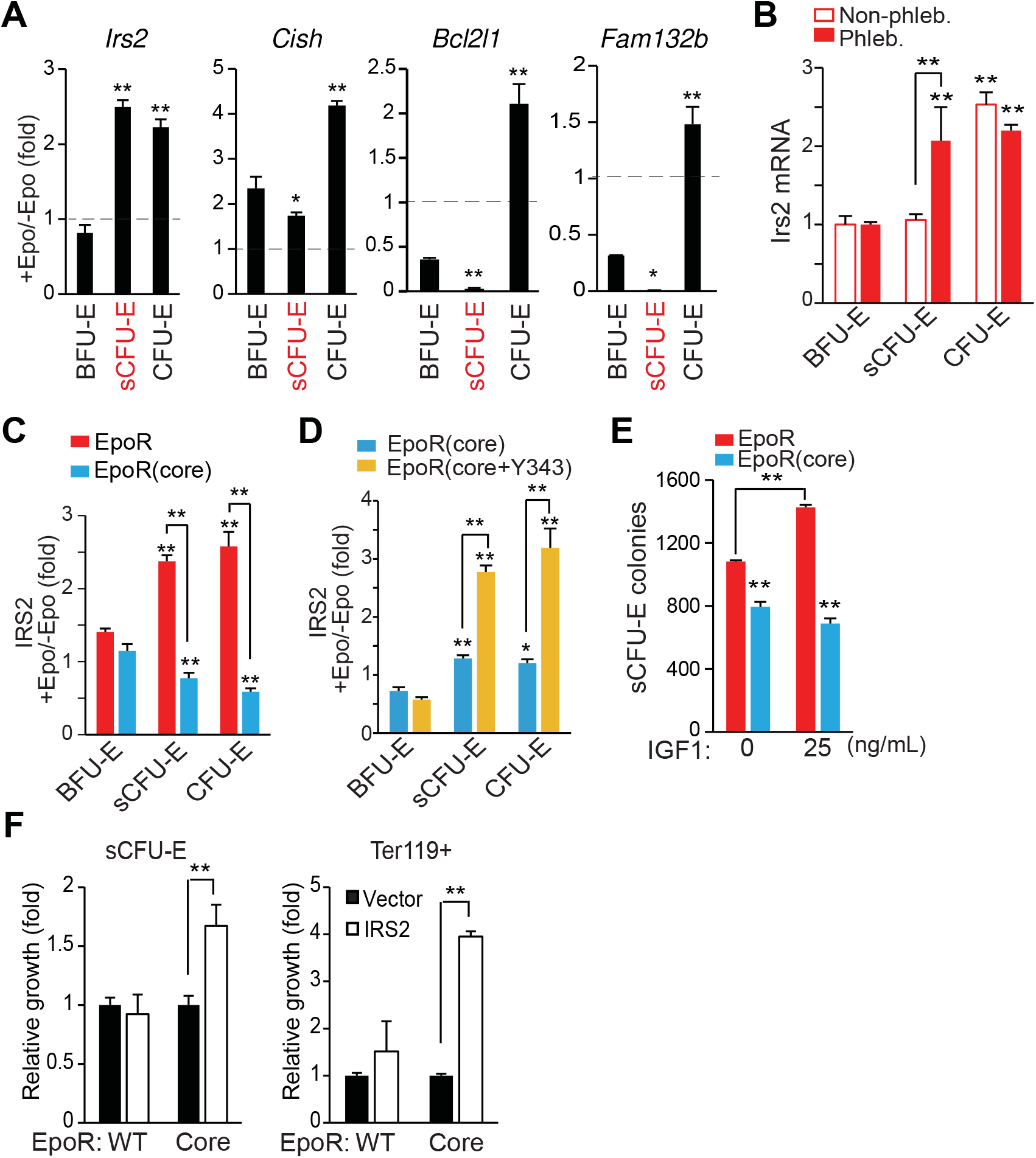
STAT5-induced IRS2 engages IGF1R signaling to promote sCFU-E growth. (A) Epo-induced expression of candidate STAT5 target genes in sorted BFU-E, sCFU-E and CFU-E by qPCR. (B) Phlebotomy induces IRS2 expression in sCFU-E. Statistically significant differences indicated on top of each bar are comparison with BFU-E, whereas significant differences between non-phlebotomized and phlebotomized conditions are specified (C) Epo-induced IRS2 expression is defective in sCFU-E and CFU-E of EpoR(core) mice. (D) Y343 rescues Epo-induced IRS2 expression in EpoR(core+Y343) sCFU-E and CFU-E. (E) IGF1 increases sCFU-E colonies in mice expressing wild-type but not EpoR(core). (F) Exogenous expression of IRS2 increases the growth of sCFU-E cells and the number of Ter119^+^ progenies generated in bone marrow cells from EpoR(core). GFP^+^ cells were gated for analyses and normalized to vector controls. *P < 0.05, **P < 0.01 analyzed by one-way or two-way ANOVA.

IRS2 is an adaptor protein that mediates signaling from both the IGF1 receptor (IGF1R) and the insulin receptor,^30^ and erythroid progenitors express higher levels of IGF1R.^25^ Consistent with a role of IGF1R and IRS2 in sCFU-E expansion, IGF1 promoted sCFU-E colony formation in wild-type but not EpoR(core) mice (Fig. 5E).

We next examined whether exogenous IRS2 expression can rescue growth of EpoR(core) sCFU-E. Lin^-^ bone marrow cells from wild-type or EpoR(core) mice were retrovirally transduced to express myc-tagged IRS2 using a bicistronic vector that also expresses GFP. These cells were first cultured in the absence of Epo to allow IRS2 expression before switching to Epo-containing media to promote erythroid cell proliferation and differentiation. Transduction efficiencies were comparable between wild-type and EpoR(core) cells (data not shown), and GFP^+^ cells were gated for analyses. Exogenous expression of IRS2 significantly expanded EpoR(core) sCFU-E and downstream Ter119^+^ progenies (Fig. 5F). Ter119^+^ progeny produced by EpoR(core) progenitors was greater in number than that produced by normal progenitors, possibly because EpoR(core), besides its inability to induce *Irs2*, is also defective in inducing negative regulators of EpoR signaling such as *Cish* (data not shown).

### Impaired sCFU-E expansion prevents erythrocytosis in JAK2(V617F)-induced myeloproliferative neoplasms

In myeloproliferative neoplasms (MPN), particularly polycythemia vera (PV), erythrocytes are overproduced as a result of abnormal and persistent high erythropoietic activity as in stress erythropoiesis. We previously showed that the EpoR, by acting as a scaffold to recruit signaling proteins, is required for the hyperactive JAK2 mutant JAK2(V617F) to drive erythrocytosis.^21^ To test whether JAK2(V617F) drives erythrocytosis via sCFU-E expansion, we employed murine bone marrow transplant models of JAK2(V617F)-driven MPN (Fig. 6A).^21^ In mice expressing normal EpoR, JAK2(V617F) drove both erythrocytosis and granulocytosis; however, in mice expressing EpoR(core), JAK2(V617F) drove granulocytosis but sCFU-E expansion was impaired and erythrocytosis was fully suppressed (Fig. 6A-C). These results were corroborated in a knock-in model of JAK2(V617F)-induced PV driven by Mx1-cre.^31^ Erythrocytosis and splenomegaly were normalized in EpoR(core)JAK2(V617F)KI mice (Fig. 6D and Supplemental Fig. 4), and the numbers of sCFU-E, CFU-E and Ter119+ cells were significantly reduced (Fig. 6E).

**Fig. 6.**
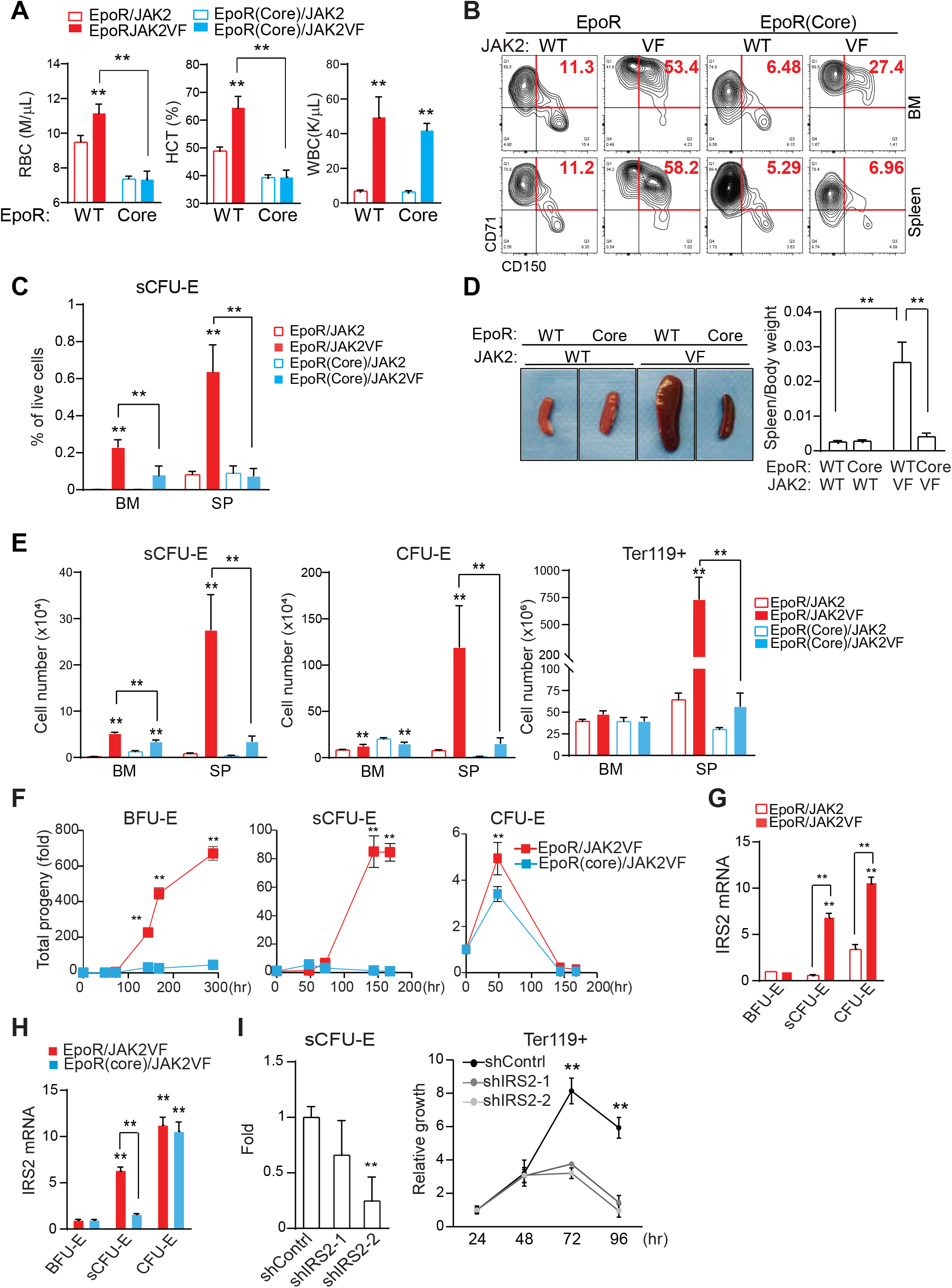
Impaired sCFU-E expansion prevents JAK2(V617F)-driven erythrocytosis in MPN. (A) Blood cell counts in transplanted mice expressing EpoR or EpoR(core) together with JAK2 or JAK2(V617F) 3 months post transplantation. RBC: red blood cell count. HCT: hematocrit. WBC: white blood cell count. (B-C) JAK2(V617F)-driven sCFU-E expansion is defective in transplant recipient mice expressing EpoR(core). Representative flow plots are shown in (B) and quantifications in (C). (D) Expression of EpoR(core) prevents JAK2(V617F)-induced splenomegaly in JAK2(V617F)KI mice. (E) The numbers of sCFU-E, CFU-E and Ter119^+^ cells are significantly reduced in EpoR(core)/JAK2(V617F)KI mice compared to EpoR/JAK2(V617F)KI mice. (F) Sorted BFU-E and sCFU-E from EpoR(core)/JAK2(V617F)KI mice fail to generate Ter119^+^ progenies *in vitro*. Cells were cultured in media with SCF but devoid of Epo. (G) In mice expressing wild-type EpoR, JAK2(V617F) increases IRS2 mRNA expression in sCFU-E and CFU-E cells. (H) IRS2 mRNA expression is significantly reduced in sCFU-E from EpoR(core)/JAK2(V617F) mice. (I) IRS2 knockdown inhibits sCFU-E and erythroid progeny growth *in vitro*. GFP^+^ cells are gated for analyses. sCFU-E fold changes are normalized to shControl, and the relative growth of Ter119^+^ cells are normalized to cell numbers at 24 hrs. *P < 0.05, **P < 0.01 analyzed by two-way ANOVA.

To directly compare the ability of JAK2(V617F) to drive constitutive signaling in cells expressing wild-type EpoR or EpoR(core), we isolated BFU-E, sCFU-E and CFU-E from EpoR/JAK2(V617F)KI and EpoR(core)/JAK2(V617F)KI mice and compared their growth *in vitro* in the absence of Epo. sCFU-E and BFU-E with a normal EpoR grew robustly in the absence of Epo, but sCFU-E and BFU-E from EpoR(core) did not (Fig. 6F). CFU-E from both animals had similar growth, indicating that signals in CFU-E and later Ter119^+^ erythroblasts are largely preserved in EpoR(core) cells (Fig. 6F). Even in cultures with Epo, BFU-E and sCFU-E expressing JAK2(V617F) with wild-type EpoR grew much better and generated more progeny than those with EpoR(core), whereas the difference in CFU-E was mild (Supplemental Fig. 5). These results show that JAK2(V617F)-dependent erythrocytosis requires induction of sCFU-E via signaling downstream of EpoR.

Consistent with the importance of IGF1R/IRS2 signaling in sCFU-E expansion, *Irs2* expression was higher in sCFU-E in mice expressing JAK2(V617F) than wild-type JAK2 (Fig. 6G). *Irs2* expression was also higher in sCFU-E from EpoR/JAK2(V617F) mice than from EpoR(core)/JAK2(V617F) mice (FIG. 6H). *Irs2* expression was normal in EpoR(core)/JAK2(V617F) CFU-E cells, indicating that there exist alternative ways to up-regulate *Irs2* in CFU-E.

To test whether IRS2 is essential for sCFU-E proliferation, we knocked down IRS2 in Lin^-^ bone marrow cells from EpoR/JAK2(V617F) mice using lentiviral shRNA vectors. After a brief culture to allow for shRNA expression, cells were cultured in low Epo media and compared for erythroid cell proliferation and differentiation. Two independent shRNAs targeting IRS2 decreased sCFU-E growth and production of downstream Ter119^+^ progeny relative to control shRNA (Fig. 6I), demonstrating that IRS2 is necessary for sCFU-E expansion in JAK2(V617F)-driven erythrocytosis.

### sCFU-E is expanded in human MPN

In humans, intermediate populations between BFU-E and CFU-E have been observed,^32-34^ and IRS2 expression increases upon erythroid differentiation in CD34+ cell cultures.^35^ We therefore tried to correlate our findings in the human setting. We examined bone marrow samples from JAK2(V617F)-positive PV patients using established immunophenotypic markers for human BFU-E and CFU-E.^26^ Samples from lymphoma and monoclonal gammopathy of undetermined significance (MGUS) that show no marrow involvement were used as controls. Within Lin^-^IL3R^-^ cells BFU-E was identified as CD34^+^CD36^-^ and CFU-E as CD34^-^CD36^+^. Stem and early hematopoietic progenitors are also included in the BFU-E gate, but their numbers are presumably low. Similar to sCFU-E observed in mice, PV samples had significantly more intermediate CD34^+^CD36^+^ cells compared to controls (Fig. 7A). PV samples also had significantly more CFU-E (Fig. 7A). Consistent with an essential role of JAK2 signaling, fewer CD34^+^CD36^+^ and CFU-E cells were observed in a PV sample from a patient treated with JAK2 inhibitors and hydroxyurea (Fig. 7A). We also established *in vitro* cultures of sorted PV BFU-E and normal controls. Similar to what was observed in murine cultures, BFU-E progressed to CD34^+^CD36^+^ cells then to CFU-E (Fig. 7B). Importantly, while the initial percentages of CD34^+^CD36^+^ cells were similar between control and PV samples (day 2), higher percentages of CD34^+^CD36^+^ cells persisted in PV cultures (Fig. 7B, Supplemental Fig. 6). Together, these results suggest that CD34^+^CD36^+^ cells are equivalent to sCFU-E observed in murine models.

**Fig. 7.**
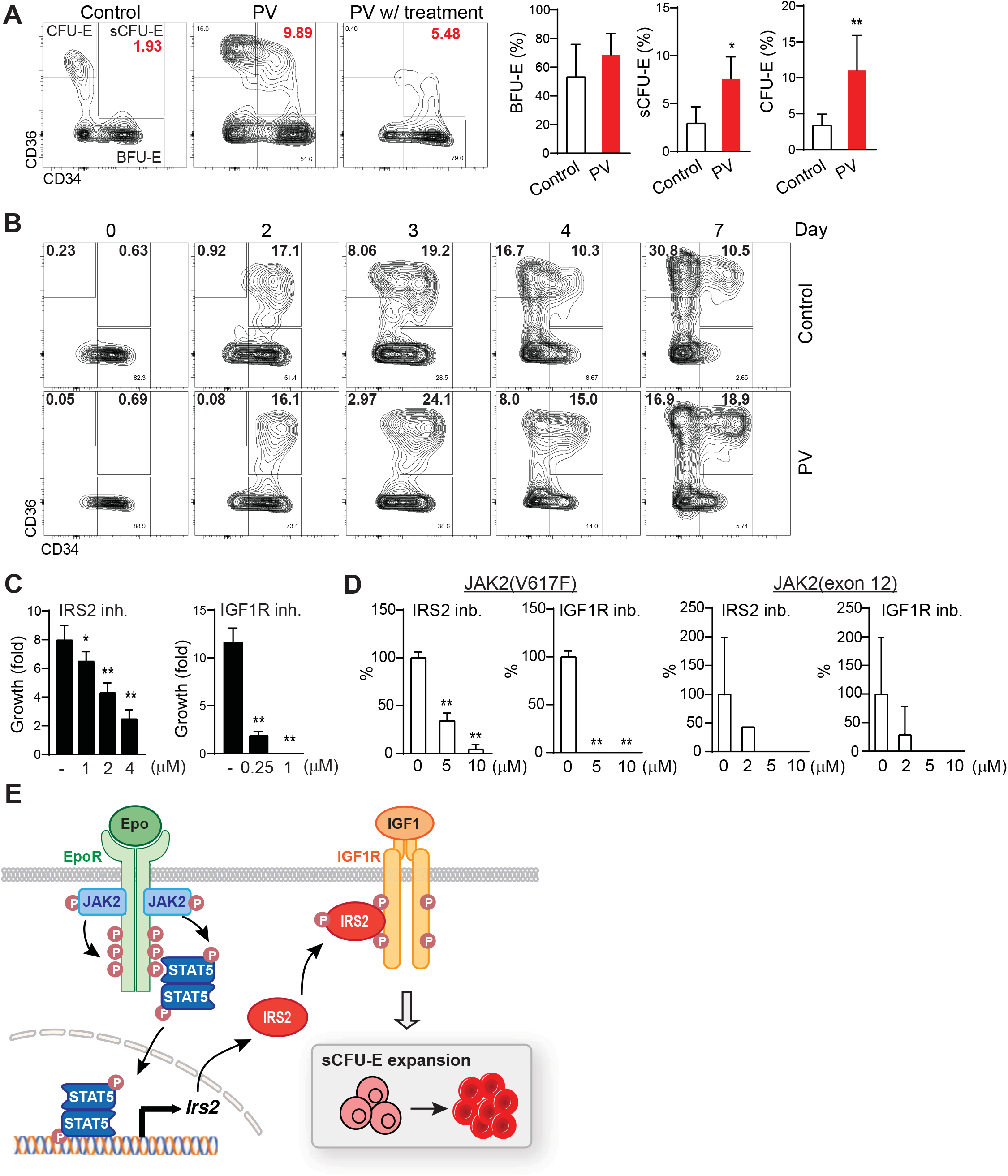
sCFU-E is expanded in human PV and inhibition of IGF1R/IRS2 signaling suppresses Epo-hypersensitive erythroid colonies. (A) Percentages of CD34^+^CD36^+^ and CD34^-^CD36^+^cells increase in PV samples. (B) *In vitro* culture of sorted BFU-E from PV or controls. Cells are examined by flow cytometry on indicated day post culture. (C) Inhibitors of IRS2 or IGF1R kinase activity reduce sorted murine sCFU-E growth *in vitro*. Data presented are 48 hours in culture (D) IRS2 and IGF1R kinase inhibitors reduce the number of erythroid colonies grown from peripheral mononuclear cells from PV patients. Cells are cultured in methylcellulose media with SCF (50ng/ml) and low Epo (0.05U/ml) and colonies are scored on day 14. (E) Current model of EpoR-IGF1R/IRS2 signaling crosstalk. *P < 0.05, **P < 0.01 analyzed by Student’s t-test or one-way ANOVA.

Ashley *et al*. recently showed that intermediate CD34^+^CD36^+^ cells could be grown from CD34^+^ cell culture *in vitro* and form CFU-E colonies.^33^ These cells can be further dissected into immature CD71^med^CD105^med^ and mature CD71^hi^CD105^hi^ subsets, and only CD71^med^CD105^med^ cells are responsive to dexamethasone.^33^ Murine sCFU-E cells also express CD71 and CD105, and their expression increases as cells differentiate from BFU-E to sCFU-E to CFU-E (Supplemental Fig. 7). Contrasting from what was observed for dexamethasone, PV CD34^+^CD36^+^ cells had increased percentages of the CD71^hi^CD105^hi^ cells, but not the CD71^hi^CD105^lo^ or CD71^lo^CD105^lo^ cells. A similar observation was made in PV mice, where the CD71^hi^CD105^hi^ subset expanded more than CD71^med^CD105^med^ subset in sCFU-E (Supplemental Fig. 8).

To examine the therapeutic potential of targeting the IGF1R/IRS2 pathway in MPN, we examined the effect of two inhibitors. NT157 causes IRS2 degradation, whereas BMS-754807 inhibits IGF1R kinase activity.^36^ Both inhibitors, in a dose-dependent manner, effectively inhibited murine sCFU-E growth (Fig. 7C) and the ability of JAK2(V617F)-positive PV mononuclear cells to form erythroid colonies under low Epo conditions (Fig. 7D). Some PV patients harbor mutations in JAK2 exon 12 instead of V617F, these inhibitors also reduced erythroid colony formation in exon 12 mutant samples (Fig. 7D).

Together, our results identify sCFU-E as a novel cell population specifically expanded in erythropoietic stress, and a synergistic crosstalk between EpoR and IGF1R, mediated by IRS2, is essential for sCFU-E expansion (Fig. 7E). Moreover, sCFU-E expansion is essential for oncogenic JAK2 mutants to drive erythrocytosis in MPN. Therapeutic targeting of sCFU-E, both positively and negatively, could be beneficial for treating anemia and MPN, respectively.

## DISCUSSION

In this study, we identify an intermediate progenitor population, hierarchically between BFU-E and CFU-E, that is specifically induced by erythropoietic stress. These sCFU-E cells are targets of rising plasma Epo, driving the recovery of peripheral RBC mass. In MPN, sCFU-E are hijacked by oncogenic JAK2 mutants to drive erythrocytosis. Molecularly, we show that signaling from the EpoR distal domain, via STAT5 induction of IRS2, engages IGF1R signaling for sCFU-E expansion. These results identify sCFU-E as targets for therapeutic interventions in anemia or MPN.

Mechanisms regulating stress erythropoiesis are best understood in the mouse spleen where it is most significantly observed. Survival of splenic Ter119^+^ erythroblasts but not their bone-marrow counterparts is regulated by Fas/FasL in stress erythropoiesis.^37^ Wnt/β-catenin signaling is not required for steady-state erythropoiesis,^38^ but is required for the proliferation of splenic stress erythroid progenitors and stress erythropoiesis.^39^ Moreover, elegant studies from the Paulson laboratory has shown that erythropoietic stress stimulates migration of short-term hematopoietic stem cells to the spleen, where they expand and differentiate into specialized “stress BFU-E”. Contrary to steady-state BFU-E, the generation of “stress BFU-E” requires splenic bone morphogenetic protein BMP4.^13,40^ Singbrant *et al*. later showed that these splenic BMP4-responsive “stress BFU-E” are enriched in Lin^-^Kit^+^CD71^lo^CD150^+^CD9^+^.^41^

In contrast to these spleen-specific mechanisms, we found that sCFU-E expansion, both in regenerative erythropoiesis and in MPN, occurs in the marrow as well as the spleen. Because human stress erythropoietic response occurs in the bone marrow, expansion of sCFUE may thus represent a more conserved mechanism. In this regard, sCFU-E may be similar to “day 3 BFU-E” cells observed after sublethal irradiation using colony assays.^42^ While normal BFU-E colonies form on day 7 post seeding, Peslak *et al*. showed that cells capable of generating BFU-E colonies on day 3 first expand in the marrow and subsequently migrate to the spleen after sublethal irradiation.^42^ “Day 3 BFU-E” cells are more mature than normal BFU-E, are Epo-responsive, and are consistent with sCFU-E being an intermediate population between immature (day 7) BFU-E and CFU-E.

The different modes and the diverse cellular entities and machineries highlight the complexity of regenerative erythropoiesis. Experimental anemia can be induced by different treatments such as phlebotomy, PHZ-induced hemolysis, irradiation, transplantation or inflammation. This mirrors the diverse clinical conditions stress erythropoiesis is involved in, including cardiac or pulmonary syndromes, anemia of multiple etiologies, chemotherapy and stem cell transplantation, or, in diseases such as MPN. It is proposed that BMP4-mediated stress erythropoiesis is specific to situations involving inflammation.^15^ Whether different response modalities are specific to different conditions, and the contribution of sCFU-E expansion in anemia induced by different means warrant further investigation.

Since their discovery in 1978,^32^ the function of intermediate progenitors between BFU-E and CFU-E has remained elusive. Our results and that of others suggest that these cells are heterogeneous, and different subpopulations are regulated specifically. For example, the more immature CD34^+^CD36^+^CD71^med^CD105^med^ cells are expanded upon glucocorticoid treatment in CD34^+^ cultures from normal but not steroid-resistant Diamond-Blackfan anemia patients.^33^ On the other hand, oncogenic EpoR/JAK2 signaling drives the expansion of the more mature CD34^+^CD36^+^CD71^hi^CD105^hi^ cells in MPN. Understanding mechanisms regulating the different subpopulations may inform erythroid biology and the development of optimal therapies for different disease states.

A role for IGF1 in expanding sCFU-E in stress erythropoiesis and erythrocytosis is consistent with prior findings that IGF1 stimulates the proliferation of erythroid progenitors.^32,43,44^ Moreover, erythroid progenitors in MPN patients are known to be hypersensitive to IGF1,^45-47^ and combinations of JAK2 and IGF1R inhibitors show therapeutic efficacy in MPN mice.^48^ Mice with poor ability to degrade the IGF1R due to deficiency of arsenite-inducible RNA-associated protein-like (AIRAPL) develop MPN.^49^ IGF1 levels decreases significantly with age, and low IGF1 levels are associated with anemia and “anemia of aging”.^50-53^ The decline in IGF1 levels may support longevity,^50^ but this may come at the cost of decreased ability to expand sCFU-E to respond to erythropoietic stress, which can exacerbate anemia in the elderly due to chronic diseases or iron deficiency.

Molecularly, the linchpin connecting EpoR and IGF1R signaling pathways in promoting sCFU-E expansion is IRS2, an adaptor protein essential for IGF1R pro-growth signaling. Epo induces IRS2 expression in sCFU-E via STAT5 activation. Both Epo and IGF1 can induce tyrosine phosphorylation on IRS2, creating docking sites for other signaling proteins. IRS2 can also undergo other post-translational modifications that alter its functions.^54-56^ IRS2 can translocate into the nucleus and forms nucleolar complexes with upstream binding factor 1, which regulates RNA polymerase I activity in rRNA synthesis,^57^ or bind to NF-kb and localize to the cyclin D promoter.^58^ Therefore, IRS2 may contribute to sCFU-E expansion and erythroid progenitor proliferation and differentiation via multiple mechanisms.

Our studies identify a new hematopoietic progenitor cell population induced to combat heightened erythroid demand in stress erythropoiesis. We also show that these cells can be hijacked to promote erythrocytosis in MPN. The lineage-restricted and stress-specific nature of sCFU-E may make them a safer and more ideal target for the treatment of both anemia and erythrocytosis. It should be noted that in MPN, simultaneous targeting of JAK2(V617F)-expressing HSC is necessary, but simultaneous targeting of expanded sCFU-E may allow for a lower HSC-directed treatment to reduce the toxicity observed in current treatment regimens.

## ACKNOWLEDGMENT

We thank Dr. Merav Socolovsky for sharing flow cytometry protocols, Drs Harvey Lodish and James Palis for EpoR(core) and EpoR(core+Y343) mice, Drs. Ben Ebert and Ann Mullally for JAK2(V617F) knockin mice, and Dr. Cheryl Lewis for assisting patient sample collection. This study was supported by research funding from the National Institute of Health (R01 HL089966 to L.J.H., R01DK111430 and R01CA230631 to J.X., R01HG011996, R01CA262781, R21CA259771 and P30CA142543 to LX, K08CA194275 to S.S.C. and P01CA108671 to R.K.R.), Rally Foundation (L.X.), the Cancer Prevention Research Institute of Texas (RP180805 and RP200103 to L.X., RR180046 to S.S.C.), American Society of hematology (Junior Faculty Scholar Award to S.S.C.), Memorial Sloan Kettering Cancer Center (P30CA008748 to R.K.R), Harold C. Simmons Comprehensive Cancer Center (pilot translational grant to L.J.H), as well as funds from Incyte, Constellation, Stemline and Zentalis to R.K.R..

## AUTHOR CONTRIBUTIONS

HH, HY, YM and LJH designed the research. HH, HY, YM, YZ, XX, HS, SCC, JX, RKR and LJH performed research studies and analyzed data. YZ, JX, XX and LX performed bioinformatics analyses. HH and LJH wrote the manuscript with input from all authors.

## References

1. Constantinescu SN, Ghaffari S, Lodish HF. The Erythropoietin Receptor: Structure, Activation and Intracellular Signal Transduction. Trends Endocrinol Metab. 1999;10(1):18–23.

2. Kuhrt D, Wojchowski DM. Emerging EPO and EPO receptor regulators and signal transducers. Blood. 2015;125(23):3536–3541.

3. Wu H, Liu X, Jaenisch R, Lodish HF. Generation of committed erythroid BFU-E and CFU-E progenitors does not require erythropoietin or the erythropoietin receptor. Cell. 1995;83(1):59–67.

4. Ihle JN. STATs: signal transducers and activators of transcription. Cell. 1996;84(3):331–334.

5. Darnell JE, Jr. STATs and gene regulation. Science. 1997;277(5332):1630–1635.

6. Richmond TD, Chohan M, Barber DL. Turning cells red: signal transduction mediated by erythropoietin. Trends Cell Biol. 2005;15(3):146–155.

7. Koury MJ, Bondurant MC. Maintenance by erythropoietin of viability and maturation of murine erythroid precursor cells. J Cell Physiol. 1988;137(1):65–74.

8. Koury MJ, Bondurant MC. Control of red cell production: the roles of programmed cell death (apoptosis) and erythropoietin. Transfusion. 1990;30(8):673–674.

9. Kelley LL, Koury MJ, Bondurant MC, Koury ST, Sawyer ST, Wickrema A. Survival or death of individual proerythroblasts results from differing erythropoietin sensitivities: a mechanism for controlled rates of erythrocyte production. Blood. 1993;82(8):2340–2352.

10. Socolovsky M, Nam H, Fleming MD, Haase VH, Brugnara C, Lodish HF. Ineffective erythropoiesis in Stat5a(-/-)5b(-/-) mice due to decreased survival of early erythroblasts. Blood. 2001;98(12):3261–3273.

11. Menon MP, Karur V, Bogacheva O, Bogachev O, Cuetara B, Wojchowski DM. Signals for stress erythropoiesis are integrated via an erythropoietin receptor-phosphotyrosine-343-Stat5 axis. J Clin Invest. 2006;116(3):683–694.

12. Zang H, Sato K, Nakajima H, McKay C, Ney PA, Ihle JN. The distal region and receptor tyrosines of the Epo receptor are non-essential for in vivo erythropoiesis. Embo J. 2001;20(12):3156–3166.

13. Paulson RF, Shi L, Wu DC. Stress erythropoiesis: new signals and new stress progenitor cells. Curr Opin Hematol. 2011;18(3):139–145.

14. Socolovsky M. Molecular insights into stress erythropoiesis. Curr Opin Hematol. 2007;14(3):215–224.

15. Paulson RF, Hariharan S, Little JA. Stress erythropoiesis: definitions and models for its study. Exp Hematol. 2020.

16. Campbell PJ, Green AR. The myeloproliferative disorders. N Engl J Med. 2006;355(23):2452–2466.

17. Mesa RA, Li CY, Ketterling RP, Schroeder GS, Knudson RA, Tefferi A. Leukemic transformation in myelofibrosis with myeloid metaplasia: a single-institution experience with 91 cases. Blood. 2005;105(3):973–977.

18. Spivak JL. Myeloproliferative Neoplasms. N Engl J Med. 2017;376(22):2168–2181.

19. Huang LJ, Shen YM, Bulut GB. Advances in understanding the pathogenesis of primary familial and congenital polycythaemia. Br J Haematol. 2010;148(6):844–852.

20. Levine RL. JAK-mutant myeloproliferative neoplasms. Curr Top Microbiol Immunol. 2012;355:119–133.

21. Yao H, Ma Y, Hong Z, et al. Activating JAK2 mutants reveal cytokine receptor coupling differences that impact outcomes in myeloproliferative neoplasm. Leukemia. 2017;31(10):2122–2131.

22. Wojchowski DM, Sathyanarayana P, Dev A. Erythropoietin receptor response circuits. Curr Opin Hematol. 2010;17(3):169–176.

23. Yao H, Ma Y, Hong Z, et al. Activating JAK2 mutants reveal cytokine receptor coupling differences that impact outcomes in myeloproliferative neoplasm. Leukemia. 2017.

24. Koulnis M, Porpiglia E, Hidalgo D, Socolovsky M. Erythropoiesis: from molecular pathways to system properties. Adv Exp Med Biol. 2014;844:37–58.

25. Tusi BK, Wolock SL, Weinreb C, et al. Population snapshots predict early haematopoietic and erythroid hierarchies. Nature. 2018.

26. Li J, Hale J, Bhagia P, et al. Isolation and transcriptome analyses of human erythroid progenitors: BFU-E and CFU-E. Blood. 2014;124(24):3636–3645.

27. Li H, Natarajan A, Ezike J, et al. Rate of Progression through a Continuum of Transit-Amplifying Progenitor Cell States Regulates Blood Cell Production. Dev Cell. 2019;49(1):118–129 e117.

28. Huang P, Zhao Y, Zhong J, et al. Putative regulators for the continuum of erythroid differentiation revealed by single-cell transcriptome of human BM and UCB cells. Proc Natl Acad Sci U S A. 2020;117(23):12868–12876.

29. Porpiglia E, Hidalgo D, Koulnis M, Tzafriri AR, Socolovsky M. Stat5 signaling specifies basal versus stress erythropoietic responses through distinct binary and graded dynamic modalities. PLoS Biol. 2012;10(8):e1001383.

30. Shaw LM. The insulin receptor substrate (IRS) proteins: at the intersection of metabolism and cancer. Cell Cycle. 2011;10(11):1750–1756.

31. Mullally A, Lane SW, Ball B, et al. Physiological Jak2V617F expression causes a lethal myeloproliferative neoplasm with differential effects on hematopoietic stem and progenitor cells. Cancer Cell. 2011;17(6):584–596.

32. Gregory CJ, Eaves AC. Three stages of erythropoietic progenitor cell differentiation distinguished by a number of physical and biologic properties. Blood. 1978;51(3):527–537.

33. Ashley RJ, Yan H, Wang N, et al. Steroid resistance in Diamond Blackfan anemia associates with p57Kip2 dysregulation in erythroid progenitors. J Clin Invest. 2020;130(4):2097–2110.

34. Yan H, Ali A, Blanc L, et al. Comprehensive phenotyping of erythropoiesis in human bone marrow: Evaluation of normal and ineffective erythropoiesis. Am J Hematol. 2021;96(9):1064–1076.

35. Machado-Neto JA, Favaro P, Lazarini M, et al. Downregulation of IRS2 in myelodysplastic syndrome: a possible role in impaired hematopoietic cell differentiation. Leuk Res. 2012;36(7):931–935.

36. Fenerich BA, Fernandes JC, Rodrigues Alves APN, et al. NT157 has antineoplastic effects and inhibits IRS1/2 and STAT3/5 in JAK2(V617F)-positive myeloproliferative neoplasm cells. Signal Transduct Target Ther. 2020;5(1):5.

37. Liu Y, Pop R, Sadegh C, Brugnara C, Haase VH, Socolovsky M. Suppression of Fas-FasL coexpression by erythropoietin mediates erythroblast expansion during the erythropoietic stress response in vivo. Blood. 2006;108(1):123–133.

38. Koch U, Wilson A, Cobas M, Kemler R, Macdonald HR, Radtke F. Simultaneous loss of beta- and gamma-catenin does not perturb hematopoiesis or lymphopoiesis. Blood. 2008;111(1):160–164.

39. Chen Y, Xiang J, Qian F, et al. Epo receptor signaling in macrophages alters the splenic niche to promote erythroid differentiation. Blood. 2020;136(2):235–246.

40. Lenox LE, Perry JM, Paulson RF. BMP4 and Madh5 regulate the erythroid response to acute anemia. Blood. 2005;105(7):2741–2748.

41. Singbrant S, Mattebo A, Sigvardsson M, Strid T, Flygare J. Prospective isolation of radiation induced erythroid stress progenitors reveals unique transcriptomic and epigenetic signatures enabling increased erythroid output. Haematologica. 2020.

42. Peslak SA, Wenger J, Bemis JC, et al. EPO-mediated expansion of late-stage erythroid progenitors in the bone marrow initiates recovery from sublethal radiation stress. Blood. 2012;120(12):2501–2511.

43. Miyagawa S, Kobayashi M, Konishi N, Sato T, Ueda K. Insulin and insulin-like growth factor I support the proliferation of erythroid progenitor cells in bone marrow through the sharing of receptors. Br J Haematol. 2000;109(3):555–562.

44. Sawada K, Krantz SB, Dessypris EN, Koury ST, Sawyer ST. Human colony-forming units-erythroid do not require accessory cells, but do require direct interaction with insulin-like growth factor I and/or insulin for erythroid development. J Clin Invest. 1989;83(5):1701–1709.

45. Correa PN, Eskinazi D, Axelrad AA. Circulating erythroid progenitors in polycythemia vera are hypersensitive to insulin-like growth factor-1 in vitro: studies in an improved serum-free medium. Blood. 1994;83(1):99–112.

46. Mirza AM, Correa PN, Axelrad AA. Increased basal and induced tyrosine phosphorylation of the insulin-like growth factor I receptor beta subunit in circulating mononuclear cells of patients with polycythemia vera. Blood. 1995;86(3):877–882.

47. Staerk J, Kallin A, Demoulin JB, Vainchenker W, Constantinescu SN. JAK1 and Tyk2 activation by the homologous polycythemia vera JAK2 V617F mutation: cross-talk with IGF1 receptor. J Biol Chem. 2005;280(51):41893–41899.

48. Basu T, Bertrand H, Karantzelis N, Grunder A, Pahl HL. Pharmacological Inhibition of Insulin Growth Factor-1 Receptor (IGF-1R) Alone or in Combination With Ruxolitinib Shows Therapeutic Efficacy in Preclinical Myeloproliferative Neoplasm Models. Hemasphere. 2021;5(5):e565.

49. Osorio FG, Soria-Valles C, Santiago-Fernandez O, et al. Loss of the proteostasis factor AIRAPL causes myeloid transformation by deregulating IGF-1 signaling. Nat Med. 2016;22(1):91–96.

50. Junnila RK, List EO, Berryman DE, Murrey JW, Kopchick JJ. The GH/IGF-1 axis in ageing and longevity. Nat Rev Endocrinol. 2013;9(6):366–376.

51. Succurro E, Arturi F, Caruso V, et al. Low insulin-like growth factor-1 levels are associated with anaemia in adult non-diabetic subjects. Thromb Haemost. 2011;105(2):365–370.

52. Marini MA, Mannino GC, Fiorentino TV, Andreozzi F, Perticone F, Sesti G. A polymorphism at IGF1 locus is associated with anemia. Oncotarget. 2017;8(20):32398–32406.

53. De Vita F, Maggio M, Lauretani F, et al. Insulin-Like Growth Factor-1 and Anemia in Older Subjects: The Inchianti Study. Endocr Pract. 2015;21(11):1211–1218.

54. Verdier F, Chretien S, Billat C, Gisselbrecht S, Lacombe C, Mayeux P. Erythropoietin induces the tyrosine phosphorylation of insulin receptor substrate-2. An alternate pathway for erythropoietin-induced phosphatidylinositol 3-kinase activation. J Biol Chem. 1997;272(42):26173–26178.

55. Sathyanarayana P, Dev A, Fang J, et al. EPO receptor circuits for primary erythroblast survival. Blood. 2008;111(11):5390–5399.

56. Peng J, He L. IRS posttranslational modifications in regulating insulin signaling. J Mol Endocrinol. 2018;60(1):R1–R8.

57. Sun H, Tu X, Prisco M, Wu A, Casiburi I, Baserga R. Insulin-like growth factor I receptor signaling and nuclear translocation of insulin receptor substrates 1 and 2. Mol Endocrinol. 2003;17(3):472–486.

58. Wu S, Zhou B, Xu L, Sun H. Interaction between nuclear insulin receptor substrate-2 and NF-kappaB in IGF-1 induces response in breast cancer cells. Oncol Rep. 2010;24(6):1541–1550.

