## Supplementary material for "Epo-IGF1R crosstalk expands stress-specific progenitors in regenerative erythropoiesis and myeloproliferative neoplasm": Supplmental

### Supplemental Methods

**Mice, phlebotomy, Epo and phenylhydrazine injection** – Mice at 3-6 month of age were used. Wild-type Balb/c and C57BL/6 mice were purchased from UT Southwestern core facility or the Jackson Laboratory. EpoR(core) and EpoR(core+Y343) mice were originally generated by Dr. James Ihle.[1] JAK2(V617F) knockin (KI) mice were kindly given by Drs. Ben Ebert and Ann Mullally.[2] All experiments involving stress erythropoiesis and bone marrow transplantation studies were performed with wild-type and EpoR(core) Balb/c mice (from Dr. Harvey Lodish) except in experiments comparing EpoR(core) and EpoR(core+Y343) mice. In this instance EpoR(core) and EpoR(core+Y343) C57BL/6 mice, both from Dr. James Palis, were used because EpoR(core+Y343) mice are only available in the C57BL/6 background. Mx1Cre:JAK2(V617F)KI mice and corresponding controls were all of C57BL/6 background. In Mx1Cre:JAK2(V617F)KI mice, polyinosinic-polycytidylic acid (pIpC; Sigma-Aldrich) at a dose of 20 μg per injection every other day for a total of four injections upon weaning to induce Jak2(V617F) expression. Phlebotomy was performed by submandibular bleeding (400μL) followed by fluid replacement with normal saline twice at 6 hrs apart on day 0. For phenylhydrazine (PHZ) treatments, mice were injected intraperitoneally with 62.5 mg/kg (low dose) or 87.5 mg/kg (high dose) PHZ on day 0 and day 1. For Epo injections, 100 U of Epoetin alpha (Amgen) was injected once subcutaneously.

**Peripheral blood analysis** – Peripheral blood collected from the submandibular vein at indicated time points was analyzed on a Hemavet automated blood counter (Drew Scientific).

**Histology and Colony-forming unit assay** – Sorted BFU-E, sCFU-E and CFU-E cells were cytospun onto coverslips and stained with May-Grünwald-Giemsa (company). Murine BFU-E and CFU-E colonies were enumerated in methylcellulose media (MethoCult M3434 and H4230,

StemCell Technologies) according to manufacturer manual. To examine the role of IGF1 in sCFU-E colony formation, homemade methylcellulose media similar to MethoCult M343 was used. Homemade methylcellulose media was supplemented with 3U/ml Epo, 50ng/ml SCF, 10ng/ml IL3, 10ng/ml IL6 but with no addition of insulin. Human erythroid colonies from PV samples were enumerated in methylcellulose media (MethoCult H4230, StemCell Technologies) with 50ng/ml SCF, 10ng/ml IL3 and 0.05U/ml Epo.

**Flow cytometry analysis of erythroid progenitors –** Bone marrow cells harvested from crushed femurs and tibia and spleen cells were suspended in phosphate-buffered saline (PBS) with 2% fetal bovine serum (FBS) and 1mM EDTA and filtered through a 70 µm filter to obtain a suspension. Cells were incubated with Fc blocker then labeled with the following cell-surface markers: biotin-conjugated lineage cocktail (Gr1, Mac1, CD3, CD4, B220, CD8, Ter119) and Streptavidin-FITC, CD117-APC-Cy7, CD55-AF647, CD105-PE, CD49f-BV421, CD150-BV650 and CD71-PE-Cy7. Lineage cocktail antibodies were purchased from Tonbo Bio and all other antibodies were purchased from BioLegend. Dead cells were excluded by 7-AAD staining.

**BrdU labeling and annexin V-binding assays** – For progenitor cells, 16 hrs post injection of BrdU (100mg/kg), BM and spleen cells were harvested, fixed and permeabilized with Cytofix/Cytoperm buffer (BD Biosciences), treated with DNaseI, and stained with Pacific Blue conjugated BrdU monoclonal antibody (Invitrogen). The percentages of cells with BrdU label were determined by flow cytometry. For precursors, cells were analyzed similarly 30 min post BrdU injection. Apoptosis was quantified by staining cells with annexin V-APC (BD Pharmingen) for 15 min and a vital dye 7-AAD and analyzed by flow cytometry.

**Inhibitor treatments –** To analyze the effects of inhibitors, cultures of sorted BFU-E or sCFU-E cells were treated with inhibitors to ERK (U0126, 10µM, Cell Signaling), STAT5 (pimozide,

10μM, Cell Signaling), AKT (LY294002, 10μM, Cell Signaling), as well as IGF1R inhibitor (BMS-754807, SelleckChem) or IRS2 inhibitor (NT157, SelleckChem) at indicated concentration. Cells were analyzed by flow cytometry at indicated timepoints.

**RNA-seq and qPCR** – Gene expression across the transcriptome in sCFU-E and CFU-E cells were compared by RNA-seq using triplicate biological samples sorted from bone marrow of day 1 phlebotomized mice. RNA-seq libraries were prepared using the illumina TruSeq RNA library prep kit with Poly(A) mRNA enrichment, and sequenced on the illumina NextSeq500 system using the high output 75bp sequencing kit. Raw reads were aligned to mouse genome (mm9) by TopHat, and differentially expressed genes were identified by Cufflinks using fold change >1.5 and FDR-adjusted $P<0.05$. For quantitative PCR, RNA was prepared from sorted BFU-E, sCFU-E and CFU-E cells using the RNeasy Plus Mini kit (QIAGEN), and subsequently reverse transcribed using SuperScript® III First-Strand Synthesis for RT-PCR (Invitrogen). cDNAs were analyzed by SsoFast™ EvaGreen® Supermix with low ROX using the indicated primer pairs.

Actin forward: 5'-CATTGCTGACAGGATGCAGAAGG-3',
Actin reverse: 5'-TGCTGGAAGGTGGACAGTGAGG-3',
Irs2 forward: 5'-CCAGTAAACGGAGGTGGCTACA-3'
Irs2 reverse: 5'-CCATAGACAGCTTGGAGCCACA-3'
Cish forward: 5'-GGATCTGCTGTGCATAGCCAAG-3'
Cish reverse: 5'-CTCGAACTAGGAATGTACCCTCC-3'
Bcl2l1 forward: 5'-GCCACCTATCTGAATGACCACC-3'
Bcl2l1 reverse: 5'-AGGAACCAGCGGTTGAAGCGC-3'
Fam132b forward: 5'-CTTCTATGCGCTTGCTGCCACT-3'
Fam132b reverse: 5'-GCATTGTGCTGGCAGAGAGACT-3'

**Retroviral expression of IRS2 and IRS2 knockdown** – To express exogenous IRS2, C-terminally myc-tagged murine IRS2 was subcloned from pBABEpuro-IRS2-Myc (Addgene) into the MSCV-IRES-GFP retroviral vector. To knockdown IRS2, retroviral shRNA vectors were generated in pWH99 that also expresses GFP.[3] The hairpin sequences targeting IRS2 are: TCTCCACTCTCTGACTATATG and CAGAACGGCCTCAACTATATC. Viral particles generated from HEK293T cells were used to infect bone marrow Lin-Kit+Sca1- cells in StemPro34 media with 4 mg/mL polybrene, 6 ng/mL IL3, 10 ng/mL IL6 and 50 ng/mL SCF by centrifugation at 1500 g, 32ºC for 90 minutes. Cells were kept in the infection media for 12 hours to allow for expression of IRS2 or IRS2 knockdown. Subsequently, cells were incubated in StemPro34 media with 2 U/ml Epo, 100 ng/ml SCF, 40 ng/ml IGF1 and 1 mM dexamethasone to allow for proliferation and differentiation of erythroid lineage cells and analyzed by flow cytometry at the indicated timepoints.

**Bone marrow transplantation** – Murine bone marrow transplantation was performed as described[4] and in accordance with University guidelines.

**Human MPN patient samples** – Bone marrow mononuclear samples were obtained from consented and deidentified PV, lymphoma and MGUS patients. Human BFU-E and CFU-E were recognized using established cell-surface markers.[5] Cells were labeled with the following antibodies: biotin-conjugated lineage cocktail (CD2, CD3, CD11b, CD14, CD15, CD16, CD19, CD56, CD19, CD16, all from BioLegend) and Streptavidin-BV711 (BD Biosciences), APC-GPA (BD Biosciences), PE594-CD41 (BioLegend), BV421-CD123 (BD Biosciences), FITC-CD34 (BioLegend), PE-CD36 (BioLegend), APCCy7-CD71(BioLegend), PECy7-CD105 (BioLegend). $Lin^{-}GPA^{-}CD41^{-}IL3R^{-}$ cells were further gated for BFU-E ($CD34^{+}$CD36-) and CFU-E ($CD34^{-}$ $CD36^{+}$). Dead cells were excluded by 7-AAD staining. Data were acquired on a Fortessa or Aria (BD Biosciences) flow cytometer and analyzed with FlowJo software (Tree Star, CA). To isolate BFU-E enriched cells from PV peripheral blood mononuclear cells, biotin-conjugated anti-CD34 antibodies were used followed by streptavidin-conjugated magnetic resin. Cells were then stained with CD34 and CD36, and $CD34^+CD36^-$ cells were isolated on an Aria (BD Biosciences). For *in vitro* cultures, sorted PV $CD34^+CD36^-$ cells were first expanded in StemSpan SFEM media with StemSpan CC100 (both from StemCell Technologies) for 3 days. Subsequently, cells were cultured in SFEM media with CC100 and Epo (3U/ml) (day 0). As controls, normal $CD34^+$ cells (Cooperative Center of Excellence in Hematology at Fred Hutch) were sorted, expanded, and cultured in parallel.

References:

1. Zang H, Sato K, Nakajima H, McKay C, Ney PA, Ihle JN. The distal region and receptor tyrosines of the Epo receptor are non-essential for in vivo erythropoiesis. Embo J. 2001;20(12):3156-3166.
2. Mullally A, Lane SW, Ball B, et al. Physiological Jak2V617F expression causes a lethal myeloproliferative neoplasm with differential effects on hematopoietic stem and progenitor cells. Cancer Cell. 2011;17(6):584-596.
3. Bulut GB, Sulahian R, Yao H, Huang LJ. Cbl ubiquitination of p85 is essential for Epo-induced EpoR endocytosis. Blood. 2013;122(24):3964-3972.
4. Yao H, Ma Y, Hong Z, et al. Activating JAK2 mutants reveal cytokine receptor coupling differences that impact outcomes in myeloproliferative neoplasm. Leukemia. 2017;31(10):2122-2131.
5. Li J, Hale J, Bhagia P, et al. Isolation and transcriptome analyses of human erythroid progenitors: BFU-E and CFU-E. Blood. 2014;124(24):3636-3645.

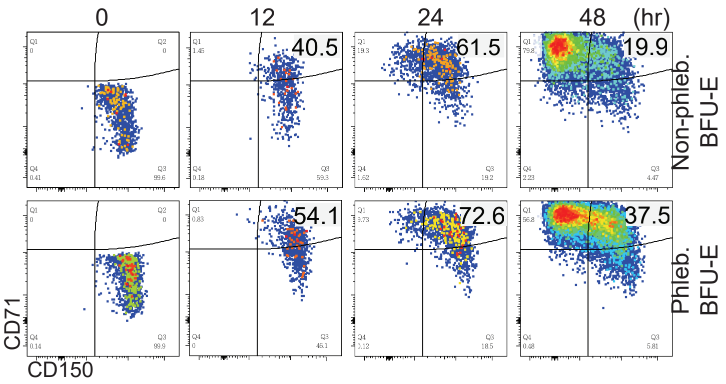


Sup. Fig. 1. *In vitro* culture of sorted BFU-E from normal or phlebotomized mice. At indicated times, cultured cells were analyzed by flow cytometry.

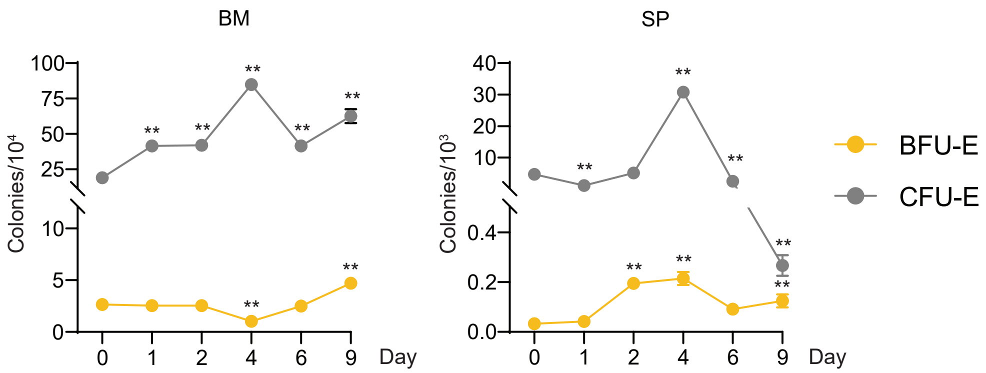


Sup. Fig. 2. CFU-E colonies significantly increase upon phlebotomy in bone marrow and spleen. Colonies were enumerated on indicated day post phlebotomy. $^*P < 0.05$, $^{**}P < 0.01$ analyzed by two-way ANOVA.

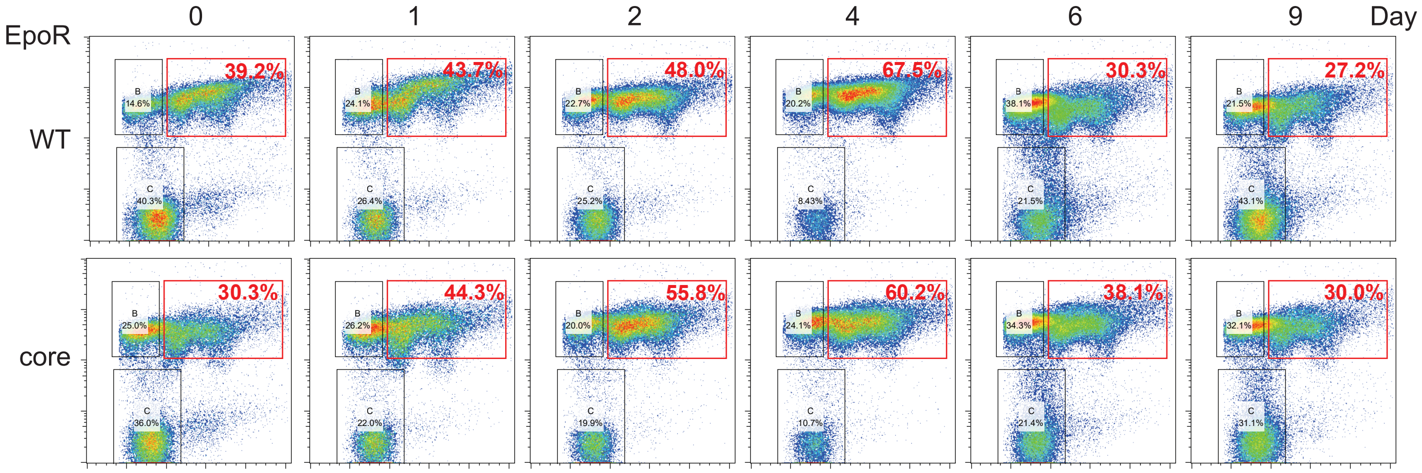


Sup. Fig. 3. Late erythroblasts expanded normally in EpoR(core) mice upon phlebotomy. Wild-type or EpoR(core) mice were phlebotomized, and at indicated days post phlebotomy, late erythroblasts in the bone marrow were analyzed by flow cytometry. Data shown are gated on Ter119$^{+}$ cells. Ter119$^{+}$FSC$^{hi}$CD71$^{+}$ cells, represent mainly basophilic erythroblasts, dramatically increased in both wild-type and EpoR(core) upon phlebotomy.

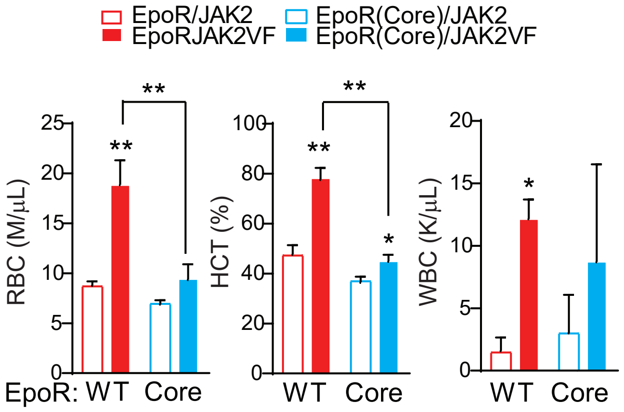


Sup. Fig. 4. Blood cell counts of JAK2(V617F) knockin mice expressing either wild-type EpoR or EpoR(core). Statistically significant differences indicated on top of each bar are comparison between cells expressing wild-type JAK2 or JAK2(V617F), whereas significant differences between cells expressing EpoR and EpoR(core) are specified. RBC: red blood cell count. HCT: hematocrit. WBC: white blood cell count. *P < 0.05, **P < 0.01 analyzed by two-way ANOVA.

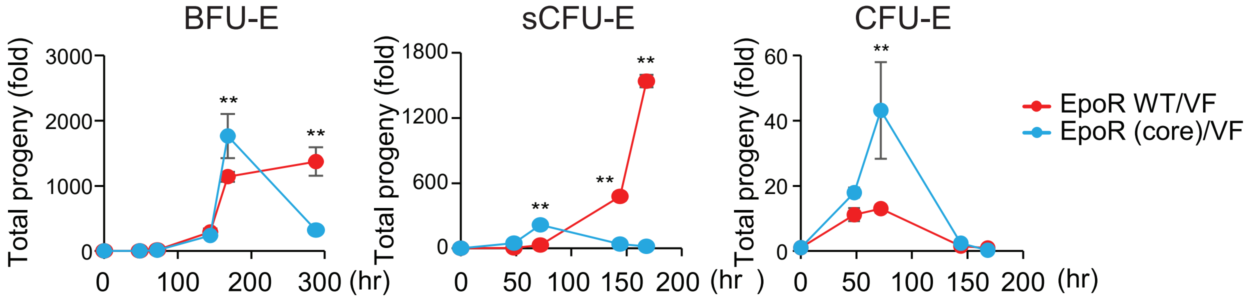


Sup. Fig. 5. Growth of sorted BFU-E, sCFU-E and CFU-E cells in the presence of Epo from EpoR/JAK2(V617F) and EpoR(core)/JAK2(V617F) mice. *P < 0.05, **P < 0.01 analyzed by two-way ANOVA.

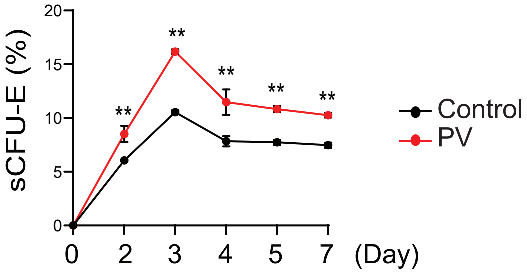


Sup. Fig. 6. Higher percentages of sCFU-E grew from PV BFU-E compared to normal controls *in vitro*. Samples were analyzed by flow cytometry at the indicated day. *P < 0.05, **P < 0.01 analyzed by two-way ANOVA.

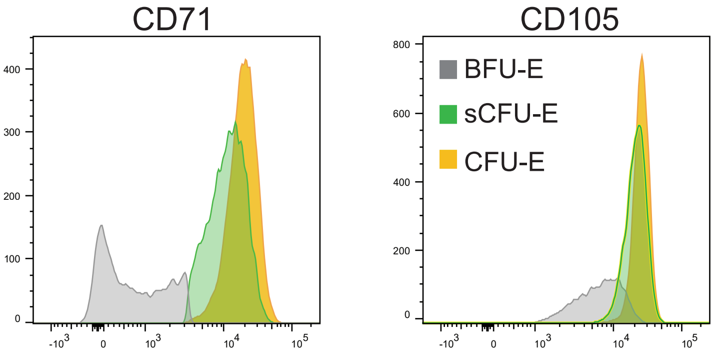


Sup. Fig. 7. CD71 and CD105 expression levels in BFU-E, sCFU-E and CFU-E from phlebotomized mice. Gated on BFU-E, sCFU-E and CFU-E cells, median fluorescence intensity for CD71 and CD105 was quantified.

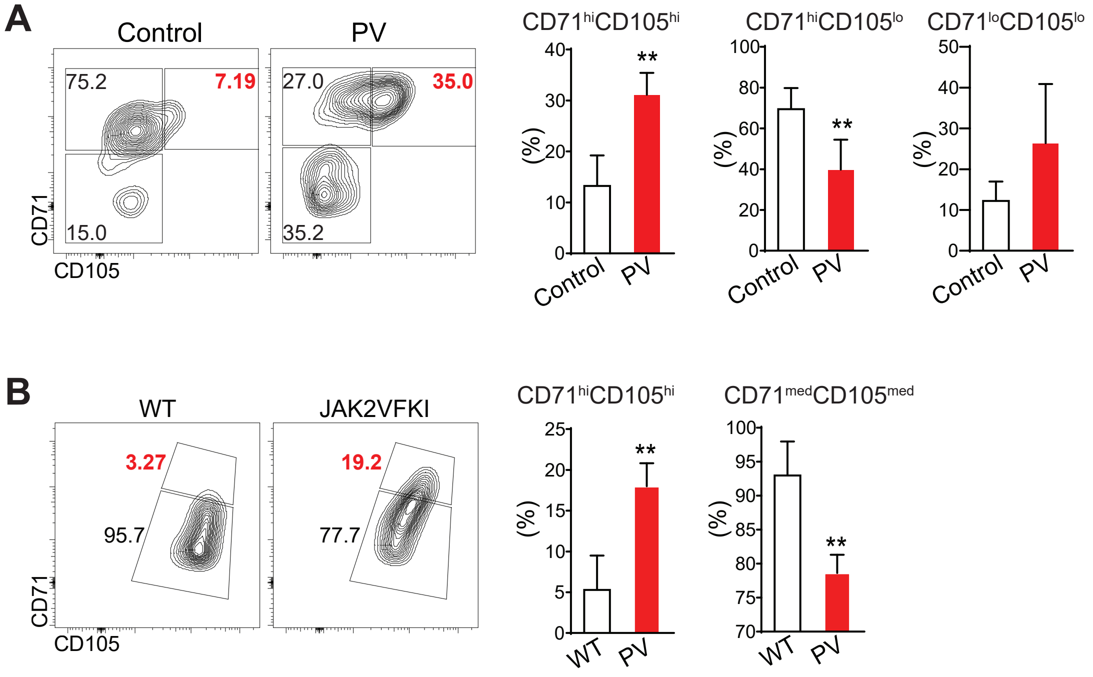


Sup. Fig. 8. Higher percentages of CD71$^{hi}$CD105$^{hi}$ cells were observed in human and mouse PV. CD34$^{+}$CD36$^{+}$ cells were gated for human (A) and sCFU-E cells were gated for murine samples (B). *P < 0.05, **P < 0.01 analyzed by Student's t-test.
